# Transcranial Focused Ultrasound Neuromodulation of the Thalamic Visual Pathway in a Large Animal Model

**DOI:** 10.1101/2021.09.21.461109

**Authors:** Morteza Mohammadjavadi, Ryan T. Ash, Pooja Gaur, Jan Kubanek, Yamil Saenz, Gary H. Glover, Gerald R. Popelka, Anthony Matthew Norcia, Kim Butts Pauly

## Abstract

**Background:** Neuromodulation of deep brain structures via transcranial ultrasound stimulation (TUS) is a promising, but still elusive approach to non-invasive treatment of brain disorders.

**Objective:** The purpose of this study was to determine whether MR-guided TUS of the lateral geniculate nucleus (LGN) can modulate visual evoked potentials (VEPs) and cortical brain oscillations in a large animal model.

**Methods:** The lateral geniculate nucleus (LGN) on one side was identified with T2-weighted MRI in sheep (all male, n=9). MR acoustic radiation force imaging (MR-ARFI) was used to confirm a tight sonication focus. Electroencephalographic (EEG) signals were recorded, and the visual evoked potential (VEP) peak-to-peak amplitude (N70 and P100) was calculated for each trial. Time-frequency spectral analysis was performed to elucidate the effect of TUS on cortical brain dynamics.

**Results:** The VEP peak-to-peak amplitude was reversibly suppressed relative to baseline during TUS. Dynamic spectral analysis demonstrated a change in cortical oscillations when TUS is paired with visual sensory input, but not when TUS is applied by itself.

**Conclusion:** TUS non-invasively delivered to LGN can neuromodulate visual activity and oscillatory dynamics in large mammalian brains.

## Introduction

In the past decade, the biophysical and cellular mechanisms of focused ultrasound neuromodulation have become better understood (Rabut *et al*., 2020). Ultrasound can activate neurons in the central nervous system either directly by opening mechanosensitive ion channels (Kubanek *et al*., 2016; Prieto *et al*., 2020; Yoo *et al*., 2020), or indirectly through changes in the composition of the lipid bilayer (Jerusalem *et al*., 2019; Sorum *et al*., 2021) inducing the flux of calcium and other ions.

Transcranial ultrasound stimulation (TUS) can be focused to noninvasively modulate neural activity in deep brain areas holding great promise for novel neurological therapies (Blackmore *et al*., 2019). However, thus far there have been few rigorous demonstrations of TUS’s impact on neural activity in an intact large animal. Sensory systems have long been used as a test bed for studying the neuromodulatory effect of TUS. The modulation of visual evoked potentials (VEPs) by ultrasound was first shown by Fry and others in 1958 (Fry *et al*., 1958). The authors showed a transient, reversible suppression of VEPs when ultrasound stimulation was directed to the LGN in craniotomized cats. However, limitations of that study included lack of detailed information on targeting, intensity, frequency and patterning of sonication, non-target control spot sonication, and sham stimulation conditions. More recent studies have demonstrated the impact of thalamic ultrasound sonication on sensory-evoked responses in larger mammals including rabbits (Yoo *et al*., 2011), pigs (Dallapiazza *et al*., 2018), sheep (Yoon *et al*., 2019), and humans (Legon et al., 2018), but a rigorous demonstration of VEP suppression with thalamic TUS in a large animal model has yet to be performed.

In addition to their role in sensory evoked responses, interactions among neuronal ensembles give rise to cortical neural oscillations observable with EEG. Several mechanisms likely contribute to synchronized oscillations in the primary visual cortex in response to visual stimuli. Cortical responses in the gamma frequency range of 30-60 Hz arise from intracortical network interactions (Georgia *et al*., 2009; Xing *et al*., 2012) as well as oscillatory input from LGN of the thalamus (Bastos *et al*., 2014). A distinction can be made between phase-locked (PL) activity and non-phase-locked (NPL) activity (Tallon-Baudry *et al*., 1996; Mike X Cohen, 2014). The PL activity typically manifests as low frequency content that is consistent with the time-locked average evoked potential whereas the NPL activity contains prominent high frequency oscillations. NPL activity provides stronger evidence for the presence of synchronized or desynchronized brain cortical oscillations than PL activity (Mike X Cohen, 2014). To the best of our knowledge, the effect of TUS on PL and NPL oscillations has not been previously investigated.

The present work sought (1) to clarify in detail how focused ultrasound targeted to the LGN modulates visual-evoked neurophysiological activity in a large mammal (sheep), and (2) to explore the broader effect of ultrasound directed to the LGN on cortical activity. Transcranial TUS was performed in lightly anesthetized sheep as a model with skull transmission properties close to human.

## Methods and Materials

### Animal Preparation and Anesthesia

All procedures were approved by the Stanford Administrative Panel on Laboratory Animal Care. Nine male sheep, 4 to 5 months old, 22 to 36 kg in body weight, were used. Four did not receive the intended TUS (one intentionally received zero TUS intensity and 3 unintentionally had transducer dislocation due to animal movement confirmed post-experiment) resulting in two experimental groups undergoing identical procedures except for the ultrasound to the LGN: an LGN TUS group (n=6, one animal underwent the experimental procedure twice with 12 days between the experiments) and an Active Sham group (n=5, one animal underwent the experimental procedure twice with 3 days between the experiments). At the end of each experiment, MR imaging was repeated to confirm the position of the transducer relative to the animal’s head, therefore the experimenters were blind to the experimental conditions for Sham or TUS groups during the experiment.

All animals were anesthetized with tiletamine and zolazepam (Telazol, Lederele Parenterals, Carolina, Puerto Rico) at 4 mg/kg, intramuscularly. The anesthesia was maintained with a combination of isoflurane delivered continuously by endo-tracheal intubation and telazol delivered by intravenous infusion. Venous and arterial catheters were placed percutaneously for drug and fluid administration and blood pressure monitoring. Lactated Ringer’s solution (Abbott Laboratories, Abbott Park, IL) was administered intravenously at approximately 10 mL/kg/hr throughout anesthesia. Isoflurane and oxygen were delivered with MRI mechanical ventilation (Omni-Vent Series D, Allied Healthcare Products, St. Louis, MO) to maintain end-tidal carbon dioxide pressure between 35 mmHg and 55 mmHg. The top of the head was shaved and treated with a depilatory cream. Pulse oximetry measurements and capnography were performed continuously during anesthesia (Expression MR400, Philips Healthcare, Vantaa, Finland).

### Imaging and Experimental Paradigm

Figure 1 illustrates the imaging and experimental paradigm. Sagittal (Panel A), coronal (Panel B) and axial (Panel C) MRI images show the TUS transducer location at control spot (i.e., away from LGN). The axial image (C) also shows the electrode locations. MR image acquisitions were performed at 3T (Signa Excite, GE Healthcare, Milwaukee, WI) using a quadrature head coil. A high resolution T2-weighted 3D FSE sequence was acquired for treatment planning with 2.5 s repetition time, 72 ms echo time, 22 cm isotropic field of view, and 256 × 192 acquisition matrix. Transducer position Panel D), transducer coupling and focal spot location were verified by MR-ARFI (Panel E) in the animals that underwent TUS. Details of the MR imaging and MR-ARFI study are elucidated in (Gaur *et al*., 2020).

**Figure 1.**
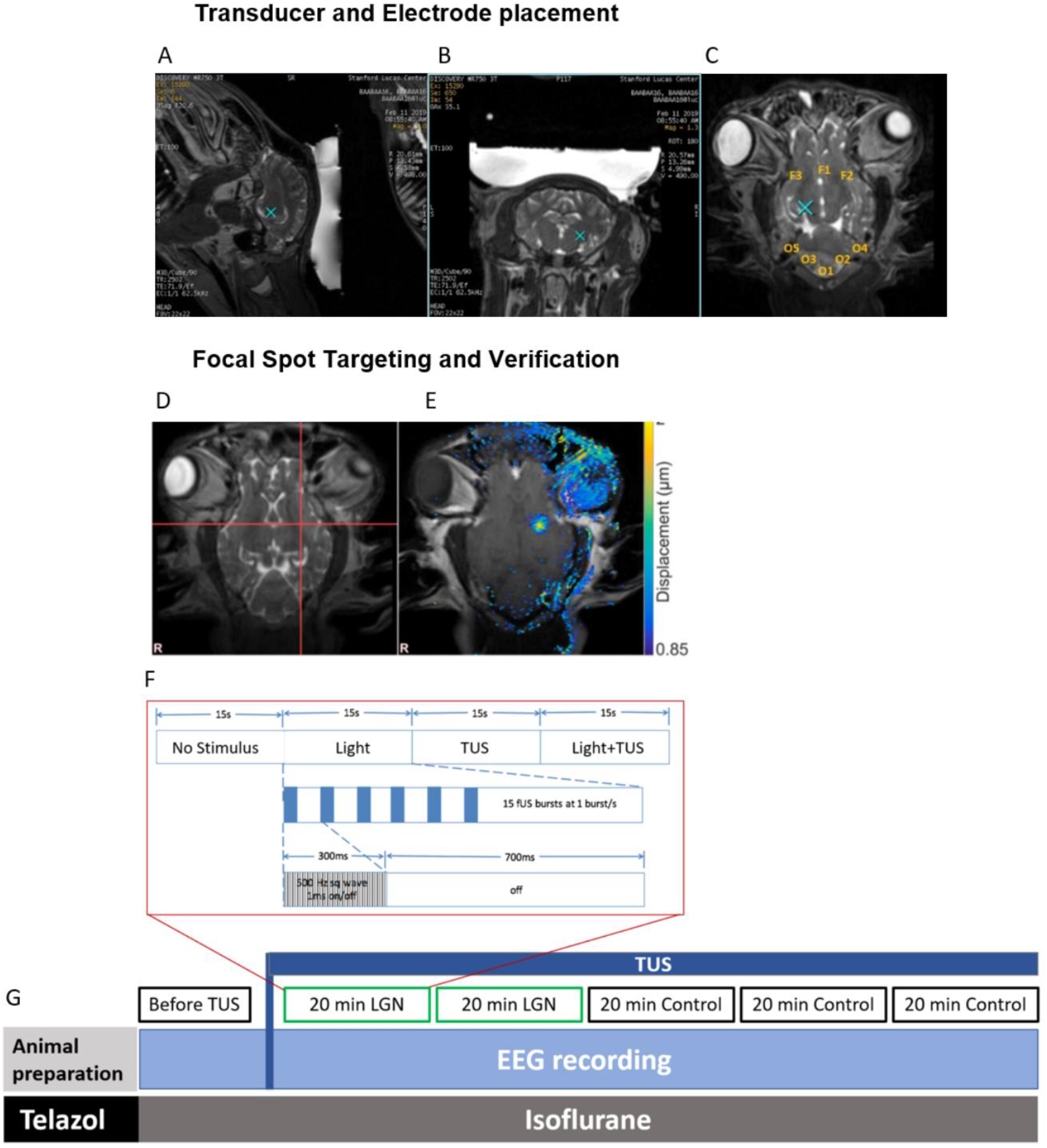
Imaging and experimental paradigm. Sagittal (**A**), coronal (**B**) and axial (**C**) MRI images with TUS target location at control spot (blue X). The axial image (**C**) also shows the eight electrode locations where each electrode was extracranially placed 10 mm under the skin (orange labels). A T2-weighted axial targeting MRI image (**D**) indicates the desired focal spot at a control location, and the MR-ARFI image (**E**) verified that the focal spot at control was at the desired location. Each 20-minute block (**F**) was split into 20 sections, each with four 15s conditions consisting of 1) no stimulus, no TUS, 2) light stimulus only, 3) TUS-only, and 4) light stimulus-plus-TUS. For each 15-second condition the repetition frequency of the stimulus was 1 flash/s. The ultrasound stimuli were 300 ms bursts of square waves of 50% duty cycle, repeated every second. The visual stimuli were 20 ms white-light flashes (binocular at 1 Hz). The block diagram of the entire experiment timeline is shown (**G**). Before applying TUS, baseline VEPs were recorded for 5 minutes. Then the 20 minute block was applied with sonication directed to either left or right LGN. This was repeated for a second 20 minute block. At that point, the sonication was directed to a control non-LGN location (10 mm anterior to LGN, medial putamen and adjacent internal capsule) for three blocks. EEG was recorded continuously during each block.

Eight platinum, monopolar, 30-gauge, 10-mm long needle electrodes were implanted subdermally on the head positioned to overlie frontal (3 electrodes) and occipital (5 electrodes) cortex (Locations illustrated in Fig. 1, Panel C). The ground electrode and reference electrode were placed on the left and right ear, respectively. A 64-channel EEG amplifier system (SynAmps RT, CompumedicsNeuroScan, Australia) with 1000 Hz sampling frequency was used. The recording system was placed in a partial Faraday cage with five of the six walls covered with aluminum foil. The light stimuli were generated with white LEDs (Linrose Electronics INC, USA) embedded at the bottom of opaque paper cups placed over the closed eyes. A 1024 element, 550 kHz focused ultrasound transducer fitted with a membrane containing chilled, degassed water (ExAblate 2100, Insightec Ltd., Haifa, Israel) was affixed to the quadrature head coil, which kept it aligned in the MR space. Degassed ultrasound gel was applied to the head for acoustic coupling and the transducer placed above the head when the quadrature MR coil was connected. During the initial MRI scanning, an additional dose of xylazine (0.1-0.2 mg/kg) was administered to minimize muscle activity before EEG signal acquisition. After that, the end-tidal volume of isoflurane was kept constant (<1%) during subsequent EEG data collection.

The visual stimuli were 20 ms white-light flashes (binocular at 1 Hz). The TUS signal was 550 kHz pulsed at 50% duty cycle for a 300 ms pulse duration. Estimates of in situ ultrasound intensity were obtained based on ex vivo fiberoptic hydrophone (Precision Acoustics, Dorset, UK) measurements of pressure transmitted through each skull cap. The power levels delivered during the study corresponded to *in situ* I_SPTA_ estimates ranging from 2.89 – 9.57 W/cm^2^ corresponding to peak negative pressure ranging from 0.55 – 1.0 MPa for neuromodulation. These values were within the FDA-approved safety limit for I_SPPA_ (< 190 W/cm^2^) but not within the I_SPTA_ (< 720 mW/cm^2^) safety limit due to the high repetition rate of sonications.

Figure 1, Panels F and G illustrate the experimental paradigm with 6 blocks of data acquisition. The first 5-minute block was a light-only stimulus condition for baseline visual activity before applying TUS. Next, there were two 20-minute blocks of LGN sonication, followed by three 20-minute blocks of non-LGN control spot sonication. The control site was approximately 10 mm anterior to the LGN and included a portion of the medial putamen and immediately adjacent internal capsule. Within each 20-minute block there were four interleaved conditions: no stimulus (15 seconds), light-only (15 1 Hz stimuli), TUS-only (15 1 Hz stimuli) and light-plus-TUS (15 1 Hz stimuli). The four conditions were cycled every minute for twenty minutes. To establish anesthesia level in relation to VEP signal-to-noise ratio, EEG was recorded during 5-minute blocks of flashing LED light only as the anesthesia lightened.

### Pre-Processing and Artifact Removal

The recorded EEG signals were processed post experiment (MATLAB 2017b, Mathworks, USA). First, a band pass filter (2-50 Hz) was applied to the continuous data, then the trials from each channel were sectioned into 1 second epochs beginning 200 ms pre-stimulus and ending 800 ms post-stimulus. Automated artifact rejection was performed with EEGLAB to remove large EEG deflections (Delorme & S Makeig, 2004). Fewer than 2% of the trials were rejected after automated pre-processing. Based on visual inspection, additional epochs clearly degraded due to high noise level or a sudden change of body movement were then manually excluded. To equalize trial number across experiments, 250 trials were considered for analysis.

Independent component analysis (ICA) was then performed (Delorme *et al*., 2004) to remove abnormal distributions in the data associated with muscle activity. Muscle activity is noticeable as oscillations of 20 to 40 Hz with relatively large amplitudes (Goncharova *et al*., 2003; Whitham *et al*., 2007). Rejected components were identified based on the following criteria: a) abnormal increase in frequency spectra around 20-40 Hz and higher, b) abnormal clustered distribution on the topographic maps that resembled scalp myogenic activity, and c) periods of abnormal power spectrum fluctuations. If the component time course showed an event related potential (ERP)-appearing signal, it was not rejected. Therefore, the ICA component removal was not automatic but a supervised process using the mentioned criteria. The main purpose of ICA analysis was to remove ultrasound artifacts from the TUS trials; therefore, ICA was not performed on Light-only conditions. Only one animal from the TUS group did not have enough EEG channels to use ICA analysis.

### Quantification of Responses

#### 1. Visual Evoked Potentials

For each electrode, block, and condition, the peak-to-peak amplitude (N70 & P100) was calculated from the averaged VEP. Because anesthesia increases the latency of the VEP peaks (Sebel *et al*., 1986; Ghita *et al*., 2013), a time window of 40 ms was defined to identify the minimum and maximum peaks for both N70 and P100, respectively (60 to100 ms time window for N70 and 90 to130 ms time window for P100). After calculating the peak-to-peak values from each electrode, the average was taken across all the electrodes. To quantify the deviation from the baseline VEP, each peak-to-peak value was normalized to the baseline condition of that animal.

#### 2. Frequency Decomposition of Brain Oscillations

Morlet wavelet convolution (Mike X Cohen, 2014) was used to analyze total power, PL and NPL spectrograms of short dynamics within post-stimulus conditions. NPL power was calculated by subtracting the average evoked-related response from each trial in the time domain and then performing a time-frequency decomposition of each trial followed by averaging across spectra. Next, PL power was calculated by subtracting the NPL power from the total power. PL activity typically contains low frequency content that is consistent with the time-locked evoked potential, whereas NPL activity includes stimulus-locked oscillations such as induced beta band (13 to 30 Hz) or gamma band (> 30 Hz) activity. A typical example is the 40 Hz visual NPL response in human (Tallon-Baudry *et al*., 1996). A range of 2 to 55 Hz with 50 frequencies was considered for calculating the spectrum. In order to highlight temporal and frequency precision, wavelet cycles in a range of 3-to 10-cycles were used. The results were converted to decibel (dB) change relative to the - 150 to 0 ms pre-stimulus baseline. Nonparametric permutation-based statistics were used to identify the significant regions of the differences between the two conditions (with no multiple comparison corrections).

## Results

### Visual evoked potentials are reversibly suppressed by sonicating thalamic LGN

Figure 2 shows VEP results for the experimental conditions. Figure 2, Panel A shows mean voltage as a function of time in the LGN TUS group at baseline (blue line, prior to sonication), during LGN sonication (red line, two LGN sonication blocks combined, light-only trials), and after LGN sonication (gray line, sonication of a non-LGN location 10-15 mm anterior to the LGN including the medial putamen and adjacent internal capsule, 3 sonication blocks combined, light-only trials). The early component of the VEP, between 50-150 ms post-stimulus-onset, can be seen clearly in the two non-LGN sonication conditions (Baseline and Control) while the LGN sonication condition clearly suppressed the VEP. The suppression recovered after sonication was retargeted to the non-LGN location. Note that the VEPs seen here were taken from the light-only condition. This rules out the possibility that the changes in VEP were due to artifacts or non-specific auditory and somatosensory-evoked EEG responses during the application of TUS. Note that the time between experimental blocks varied from experiment to experiment, so a direct quantification of the duration of the long term suppression was not possible with our measurements. Figure 2 B shows the same data for Active Sham animals in which the sonication target was determined post-experiment to be an off-target location. Figure 2 C shows VEP Peak-to-peak amplitude measurements in the LGN TUS group for the six experimental conditions including mean (solid black), +/-SEM (gray band) and individual data (N=6, colored lines). Figure 2, Panel D shows the same results as in Figure 2 C but for the Active Sham group. In the LGN TUS group, sonication caused a significant decrease in response amplitude that returned to baseline on average (Figure 2 C), while Active Sham stimulation had no reliable impact on VEP amplitude (Figure 2D). Figure 2E and F plots the mean +/- SEM VEP peak-to-peak amplitudes for the Baseline, LGN2 and Non-LGN Control blocks in the LGN TUS group (Panel E, blue bars) and the Active Sham group (Panel F, grey bars). The mean VEP amplitude decreased by almost 50% during LGN sonication compared to baseline and then recovered to baseline-like levels during subsequent control sonication blocks (Fig. 2E,baseline vs. LGN sonication: ***p = 0.0007; LGN sonication vs. control sonication: **p = 0.006, paired one-tailed t-test). Sham off-target sonication, in contrast, had no significant impact on VEP amplitude (Fig. 2F, all comparisons n.s.).

**Figure 2.**
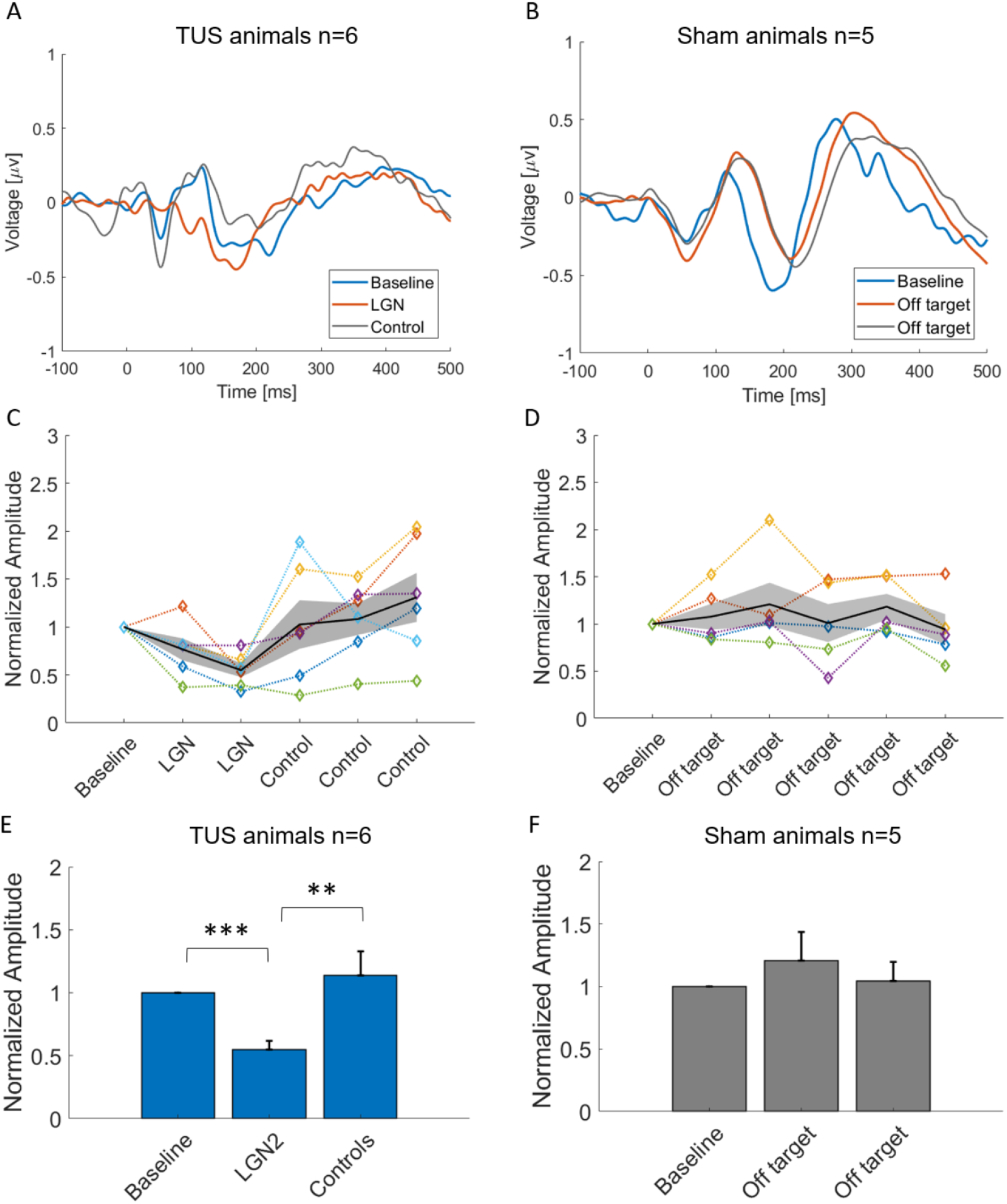
Reversible suppression of VEPs by LGN ultrasound sonication. (**A**) Grand-average VEP responses to 20-ms binocular light-flash visual stimuli before (blue line, Baseline), during (red line, LGN), and after (gray line, Control stimulation site) sonication of the LGN. Traces were normalized to the maximal response at baseline per animal, then averaged across experiments (N=6 experiments from 5 animals). For LGN and control VEPs, data were combined across blocks (2 for LGN and 3 for control). Only responses from the light-only condition (i.e. trials without TUS) were included in the analysis. Traces were smoothed with a 15-point (15 ms) moving averages filter for illustration purposes only. For a subset of animals, the EEG trace from -10 to +40 ms was removed and interpolated to eliminate a light-flash-related artifact, for illustration purposes only. (**B**) Data presented as in Panel A, for the Active Sham condition (in which ultrasound sonication was determined to be off-target post-experiment, N=4) and a no-FUS control animal (N=1,total N=5). (**C**) Peak-to-peak VEP amplitude measurements for individual experiments (colored lines) and Mean +/- SEM across experiments (black line), for the six experimental blocks (Baseline, two LGN sonications and three non-LGN sonications), for LGN-sonication animals. (**D**) Same as C, for Sham condition animals, across the six experimental blocks (Baseline and five off-target sonications). (**E**) Mean (+/- SEM) VEP peak-to-peak amplitude at baseline, LGN2, and pooled control sonication blocks for LGN-TUS (blue bars). The mean VEP amplitude during the second LGN sonication was significantly decreased compared to Sham VEP amplitudes (**p = 0.006, ***p = 0.0007, paired one-tailed t-test, N=6 LGN, N=5 Sham). (**F**) Same as E, for Sham (grey bars) animals. All comparisons nonsignificant.

We next analyzed the immediate and lasting impact of TUS as well as the direct EEG responses to TUS without any concomitant light stimulus. Figure 3 illustrates mean VEP traces (first row) and mean response amplitudes (second row) for the Light-only (column A), Light-plus-TUS (column B) and TUS-only conditions (column C) for the LGN (red) and non-LGN (gray) sonication conditions. These data reveal 3 essential points: 1) LGN sonication causes a VEP response suppression that is observable both during LGN sonication, and in light-only trials between sonication trials. The impact of LGN sonication was similar in the light-only and light-plus-TUS conditions, suggesting that the major effect of sonication on VEPs is not due to an immediate impact of sonication but rather reflects a lasting suppression that outlasts the period of sonication. This is further supported by the fact that LGN sonication’s impact reaches significance in the Light-only condition (p=0.004) but not the Light-plus-TUS condition (p=0.07); 2) there are no observable differences in VEP waveform for the Light-only condition and Light-plus-TUS condition for the LGN or non-LGN sonication conditions, indicating that TUS by itself does not dramatically impact EEG recording quality; and 3) The EEG response in the TUS only condition is minimal. Aside from a single outlier in the LGN condition, there was no clearly observable VEP response, indicating that there is little-to-no nonspecific auditory or somatosensory-evoked responses due to application of TUS.

**Figure 3.**
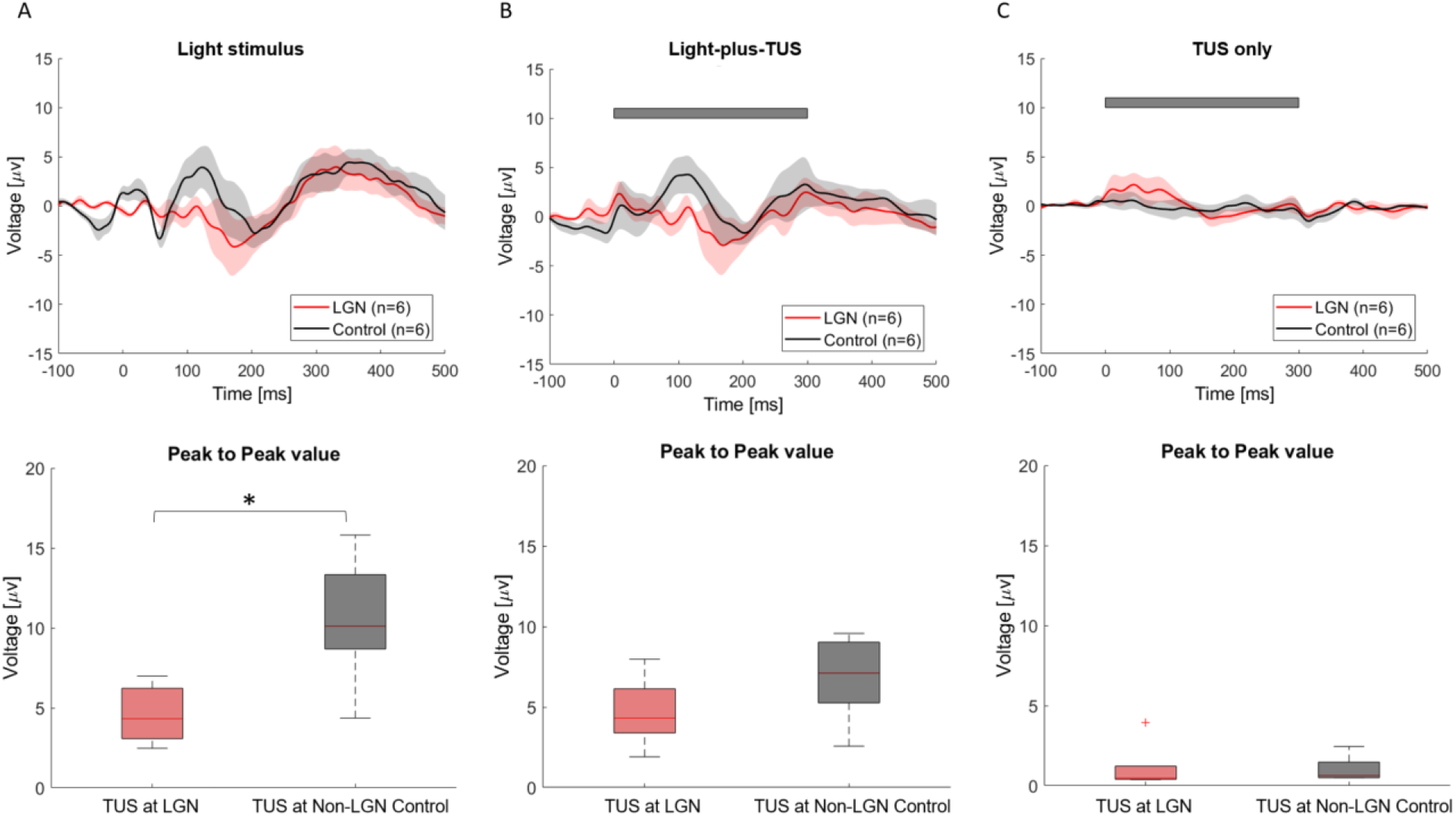
The impact of LGN TUS on VEPs outlasts the duration of sonication. Effect of LGN (red) and non-LGN control (gray, 10-15 mm anterior of LGN) sonication on VEPs for the three interleaved stimulus conditions (Light-only, Light-plus-TUS and TUS-only). (**A**) Top row: Mean +/- SEM VEP waveforms in the Light-only condition for LGN (red line, N=6) and non-LGN control (gray line, N=5) sonication. Responses were smoothed (15-point moving averages filter) for illustration purposes. Bottom row: Boxplots of VEP Peak-to-peak amplitude for LGN and non-LGN control sonication, light-only trials. (*p = 0.004, one-tailed unpaired t-test). Box depicts interquartile range, and dashed error bars depict full range of values. (**B**) Data presented as in A, but for the light-plus-TUS condition. Gray bar illustrates time course of TUS relative to visual stimulus onset. Difference between LGN and non-LGN: p=0.07. (**C**) Data presented as in A, for the TUS-only condition (i.e., no visual stimulus). Note the minimal overall response, indicative of little-to-no nonspecific auditory or somatosensory responses to transducer vibration. The transient ∼100 ms increase in voltage at TUS onset with LGN stimulation, is primarily due to responses in a single animal (see outlier dot in bottom row), indicative potentially of a subtle depolarizing effect of LGN-TUS in this animal, and/or an EEG artifact of TUS.

Given that the visual stimulus was binocular, and the LGN sonication was unilateral, visual cortical responses should decrease more ipsilateral to sonication compared to contralateral sonication. We attempted to assess the laterality of the responses by subtracting the contralateral VEPs from the ipsilateral VEPs per animal. A strong trend to a lateralized effect of LGN-TUS was observable (data not shown), but due to malfunction and disconnection of a subset of electrodes in 1 out of 6 LGN experiments and 1 out of 5 Active Sham experiments, our results were not sufficiently powered to confirm lateralization of response suppression.

### Modulation of light-elicited brain oscillations by applying ultrasound to LGN

The effect of ultrasound applied to LGN on brain oscillations is shown in Figure 4 with average spectrograms (N=5) obtained during the light-only condition (Panel A, top row) and during the light-plus-TUS condition (Panel B, bottom row). Comparing Panel A with Panel B shows that application of TUS at LGN induces NPL oscillations at upper beta and gamma-band frequencies at approximately 90 and 280 ms post-stimulus. There is also a slight PL activation in lower frequencies spread across theta and alpha bands (4-10 Hz). Increased lower band alpha power with TUS suggests that there may be less alpha desynchronization due to suppressed corticothalamic input (Pfurtscheller, 1992). The modulation of PL power is restricted to frequencies below 20 Hz.

**Figure 4.**
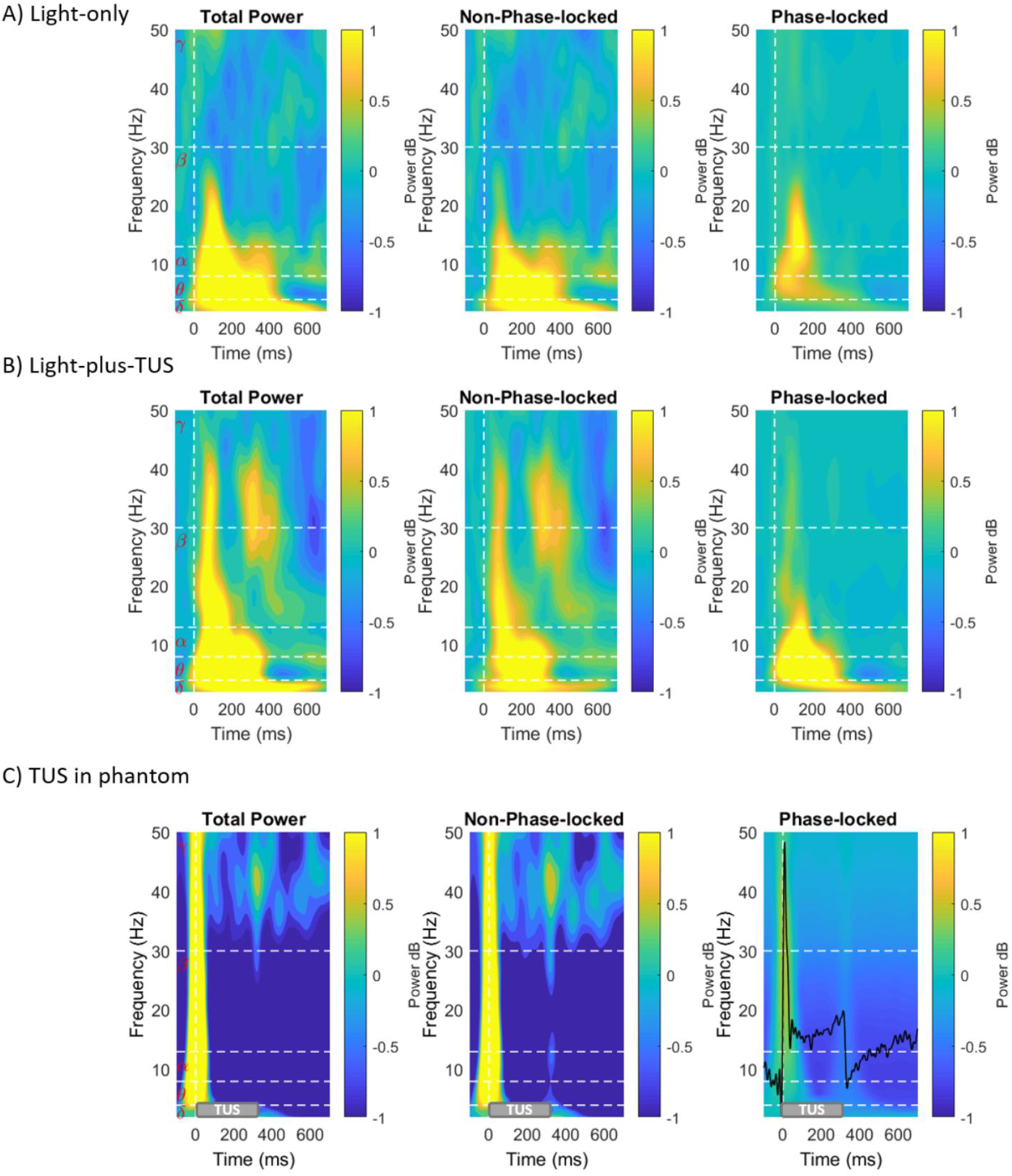
Effect of LGN TUS on Non-phase-locked and Phase-locked visual evoked cortical oscillations and EEG spectra of phantom experiments. Average (N=5) EEG power spectra for Total Power, NPL Power, and PL Power (Columns) for two experimental conditions (Rows) (A) Light-only and (B) Light and TUS simultaneously; and (C) phantom experiments. Comparing Panel A with B shows the application of TUS at the LGN induces oscillations in the occipital cortex for the upper beta and gamma-band frequencies at approximately 90 and 280 ms post-stimulus. There is also a light PL activation in lower frequencies spread across theta and alpha bands. (C) The phantom was sonicated with 300 ms bursts at 1 Hz, while recording electrical activity (N=5). The time domain signal plotted on top of the spectrum (black line); the gray bar shows 300 ms TUS duration. The frequency regions that differ significantly are shown in Figure 5.

Activation of the US transducer can induce an electromagnetic interference on the electrodes. The properties of the ultrasound artifact were quantified with experiments in which a rectangular block of tofu was used as a phantom with 5 EEG electrodes inserted and the ultrasound transducer placed directly on the surface. The frequency spectra of the EEG are shown in Figure 4C with the artifact shown as a signal commensurate with the TUS signal. When the transducer was removed from the surface and positioned at increasing distances, the magnitude of the artifact decreased as the transducer distance increased. This suggests that the source of the artifact was due to electromagnetic coupling of the transducer to the electrodes.

Figure 5 illustrates differences between spectra in Figure 5A and B. The differences are shown as spectral difference maps (upper row) for two conditions, light-only and light-plus-TUS at LGN (Column 5A) and light-only and light-plus-TUS at the non-LGN control location (Column 5B). Each of the two conditions contains difference maps for NPL and PL analyses. A nonparametric permutation-test statistic was performed (not corrected for multiple comparisons) on the difference maps between the spectrograms with differences that were statistically significant (lower row). During TUS of the LGN, significant differences in the NPL analysis were seen in the upper beta and low gamma frequency bands (30-50 Hz) at approximately 90 and 300 ms post-sonication. By contrast, in the PL analysis, significant difference was seen in the lower frequency theta and alpha bands between 90 to 300 ms. During the non-LGN control (a portion of the medial putamen and immediately adjacent internal capsule) TUS, a statistically significant reduction in the theta and alpha activity of the NPL response was observed, with no clear change in the PL response. There was no significant difference between the spectra of LGN and Non-LGN control sonication during TUS-only condition. A nonparametric statistical test was performed on the difference map between the phantom data (see Figure 4C) and the light-only baseline condition. The black contours in Figure 5A (lower row) show the significant regions due to TUS in phantom (tofu). There is a slight overlap in low frequency (< 10 Hz) content of PL activity during the application of TUS, whereas there is no overlap in the significant regions of NPL activity, indicating that the observed effects of TUS on gamma-beta frequency powers are unlikely to be due to temporal properties of the transducer’s electromagnetic artifact.

**Figure 5.**
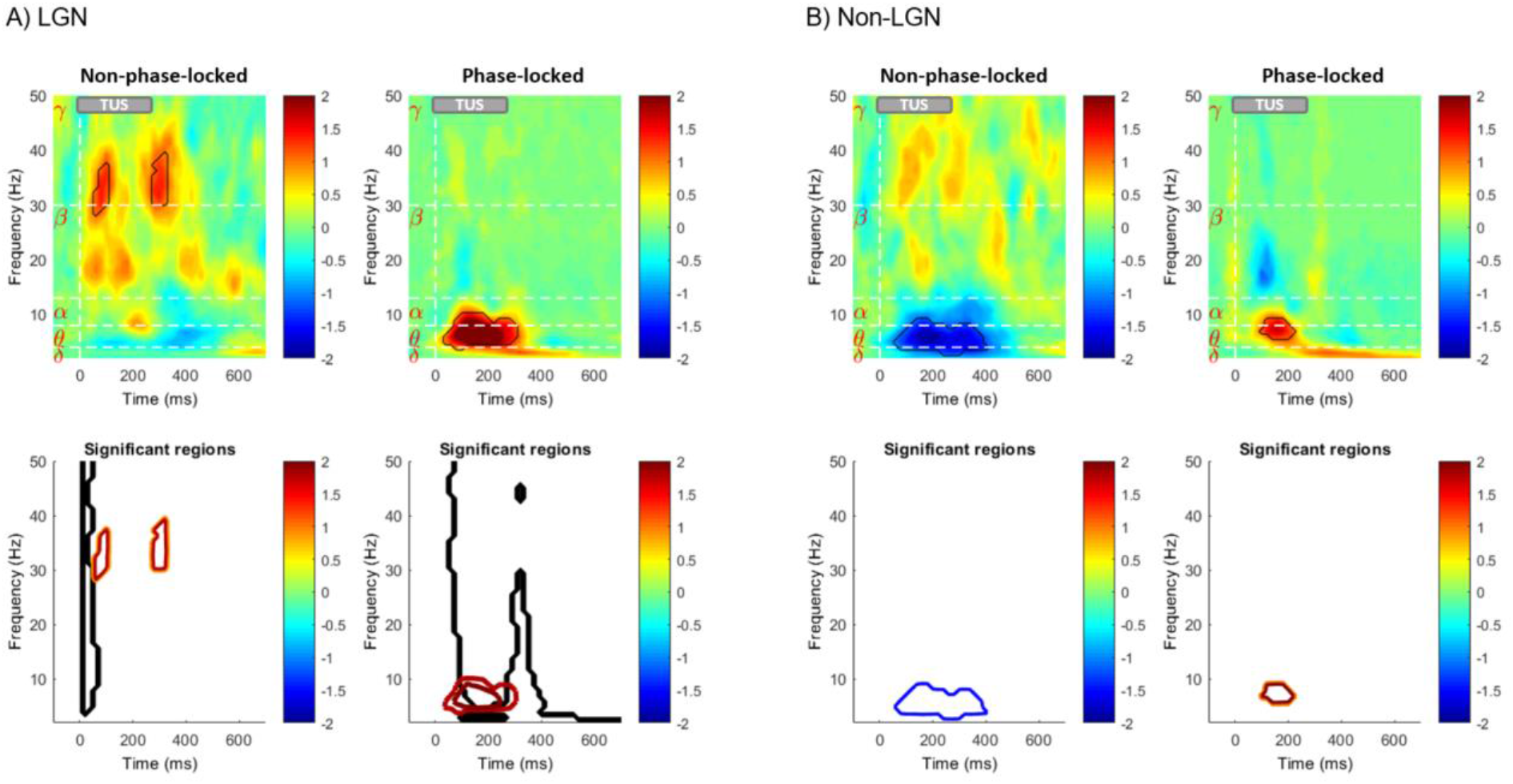
Immediate impact of LGN TUS on Non-phase-locked and Phase-locked visual evoked cortical oscillations. Differences between spectra in Figure 4. The differences are shown as spectral difference maps (upper row) for two conditions, light only and light + TUS at LGN (Column 6A) and light only and light-plus-TUS at the non-LGN control location (Column 6B). Red coloring indicates areas where TUS increased visual-evoked power, while blue coloring indicates areas where TUS decreased visual-evoked power. Each of the two conditions contains difference maps for NPL and PL analyses. For each, the contour of significant regions based on nonparametric permutation-based statistics are shown in the bottom row of each panel (p<0.05). Significant regions of spectral difference maps between tofu and light-only baseline condition are shown in black in lower row of column A. The black significant regions of Panel A are shown for only positive values.

## Discussion

We demonstrated that TUS targeted at thalamic LGN reversibly suppresses VEPs in a large animal. Our VEP analysis corroborates the results of Fry et al. in which they showed a reversible reduction in the magnitude of the light elicited VEPs by over 50% with unilateral LGN sonication. The suppression was most consistent at the second LGN block suggestive of a cumulative impact of sonication, and lasted past the LGN sonication blocks in a subset of animals, indicating an enduring effect of sonication. Comparisons of EEG responses to the light-only, the light and TUS combined, and TUS-only additionally revealed that in our preparation TUS generates a minimal if any EEG response by itself, and most or all of its impact on VEP suppression is observable in interleaved light-only trials in which there is no co-occurring TUS. This rules out a number of potential TUS-related confounds, including electromagnetic TUS artifacts, auditory-evoked or somatosensory-evoked EEG responses to vibrations or sounds made by the transducer, as potential explanations for our results. We did not see consistent evoked responses elicited by application of TUS-only to the LGN over a range of I_SPTA_ intensity from 3 to 10 W/cm2. This is in line with the majority of large animal studies which primarily show a suppressive impact of TUS. The ultrasound parameters used in this study are within the FDA-approved safety range for spatial pulse peak average (I_SPPA_) but not the spatial pulse temporal average (I_SPTA_) intensity. A number of studies including those from our lab have suggested that the FDA guidelines particularly for I_SPTA_ are very conservative. A thorough histological examination of the brains of the sheep used in this study found no evidence of any microscopic tissue injury or damage (Gaur *et al*., 2020), bolstering the clinical relevance of these findings.

On its face, a greater-than-50% decrease in the response to unilateral LGN sonication is larger than expected for a binocular visual stimulus, given that a single LGN should transmit at most 50% of the visual-evoked activity, and therefore a 100% inhibition should lead to an at-most 50% suppression in visual cortex. There are some speculations to explain this. First it is possible that sonication of the LGN changes the temporal dynamics of the volley to cortex, which in turn could affect the magnitude of its response. Second other thalamic circuitry pathways particularly the reticular nucleus could be activated by TUS during the stimulation of LGN leading to more widespread inhibitory effects.

### Modulation of cortical activity through the lens of spectral analysis

In the visual system, the LGN and primary visual cortex (V1) interact in a cortico-geniculate feedback loop via the alpha-band (8 -14 Hz) and through the beta-band (15 - 30) in geniculate-cortical feedforward processing, whereas gamma-band oscillation (>30 Hz) is largely a property of cortex (Georgia *et al*., 2009; Bastos *et al*., 2014). The origin and the functionality of these oscillations in primary visual cortex are not completely understood. To compound this further, an important body of literature exists linking the effect of volatile anesthetics to the suppression of EEG high frequency bands (Miller *et al*., 2004; Purdon *et al*., 2015; Akeju *et al*., 2016). In our study, taking into account the influence of anesthesia on high frequency oscillations, we show that the upper beta-band and gamma-band activity are increased by ultrasound neuromodulation of the thalamic visual pathway. The results from simultaneous application of TUS to LGN and visual stimulation suggest that TUS of LGN amplifies visual-evoked NPL brain oscillations that occur with time and frequency characteristics analogous to the cortical oscillations elicited by visual responses (Tallon-Baudry *et al*., 1996; Bastos *et al*., 2014). This suggests that high frequency NPL cortical oscillations are modulated by oscillatory input from the LGN. Legon et al (Legon *et al*., 2018) showed a time-locked gamma activity reduction around 100 ms and 200-300 ms post-stimulus-onset when sonicating the ventro-posterior lateral (VPL) of thalamus in human. In our study, the spectral results show an increase in the gamma-band activity occurring around the same time at 100 ms and 300 ms. Although the study design, species and targeted area were different in these two studies, the similarity in time of the effect and the frequency content of the oscillations caused by TUS targeted to the adjacent thalamic nuclei of the brain is worth further investigation in the future studies.

Initially there was a concern about the confounding effect of TUS artifact on our spectral findings. Although the ICA was efficient in eliminating the artifact, as an additional control we compared the results from a phantom experiment in tofu with the results of the sheep study. The significant regions did not overlap in NPL activity, however there might be a slight overlap in slow oscillations in the PL regions. This suggests that the effects of TUS on EEG band power are not due to the TUS artifact but rather due to changes in neural activity. We also note that our main findings on VEP suppression are not confounded by sonication artifacts, as these analyses were performed on light-only (i.e. visual stimulus only) trials within the sonication blocks, in which light-only, light-plus-TUS, and TUS only trials were interleaved.

Major limitations of the current study include 1) the timing between sonication of LGN and Non-LGN control experimental blocks was variable across animals, preventing an assessment of the time-course of suppression; 2) the visual stimulus was full-field binocular not hemi-retinal and some EEG electrodes failed in a subset of animals, preventing a well-powered assessment of the laterality of the sonication’s effect; 3) the binocular light flash stimulus did not generate robust single-trial VEPs in all animals, so responses had to be averaged across multiple trials to see any effect, preventing an analysis of the build-up of sonication effects in initial trials; 4) the control sonication position was not well-defined and included both the medial putamen and a portion of the internal capsule, so the interesting effect of non-LGN sonication on cortical oscillations (Figure 5) is difficult to interpret.

Future work should apply different measurement techniques, including PET, fMRI, and implanted electrodes to investigate the effect of TUS on thalamus in other brain regions. Our results lend further support to the growing consensus that transcranial ultrasound neuromodulation will in the future enable noninvasive deep brain stimulation therapy for a range of neuropsychiatric illnesses.

## Conclusion

Unilateral TUS of LGN can reversibly suppress binocular VEPs in sheep. This suppression returns to baseline after termination of sonication. Light-induced oscillations can be modulated by presenting a visual stimulus simultaneous with TUS to LGN.

## Acknowledgments

The authors would like to thank Karla and Kevin Epperson for help with MRI and preparation, Samuel Baker, Cholawat Pacharinsak and Benjamin Franco for help with sheep. This work was supported by NIH R01 MH111825, NIH R01 EB019005 and NIH R01 NS112152.

## Conflicts of interest

No potential conflict of interest was reported by the authors.

## Notes

### Competing Interest Statement

The authors have declared no competing interest.

